# Visible Light Plasmon Excitation of Silver Nanoparticles Against Antibiotic-Resistant *Pseudomonas aeruginosa*

**DOI:** 10.1101/2020.01.10.902676

**Authors:** Rafael T. P. da Silva, Marcos V. Petri, Estela Y. Valencia, Pedro H. C. Camargo, Susana I. C. de Torresi, Beny Spira

**Author notes:** Address correspondence to Beny Spira. Rafael T. P. da Silva, Marcos V. Petri and Estela Y. Valencia contributed equally to this work. Author order was determined by consensual agreement.

## Abstract

The interaction of metallic nanoparticles with light excites a local surface plasmon resonance (LSPR). This phenomenon enables the transfer of hot electrons to substrates that release Reactive Oxygen Species (ROS). In this context, the present study was aimed at enhancing the antibacterial effect of citrate-covered silver nanoparticles (AgNPs), which already possess excellent antimicrobial properties, via LSPR excitation with visible LED against *Pseudomonas aeruginosa*, one of the most refractory organisms to antibiotic treatment. The Minimum Inhibitory Concentration (MIC) of AgNPs was 10 μg/ml under dark conditions and 5 μg/ml under light conditions. The combination of light and AgNPs led to 100% cell death after 60 minutes. Quantification of ROS via flow cytometry showed that LSPR stimulated AgNPs increased intracellular ROS concentration by 4.8-fold, suggesting that light-exposed AgNPs caused cell death via ROS production. Light exposition caused a small release of silver ions (0.4%) reaching a maximum after 6 hours. This indicates that silver ions play at most a secondary role in *P. aeruginosa* death. Overall, the results presented here show that LSPR generation from AgNPs by visible light enhances the antimicrobial activity of silver nanoparticles and can be an alternative for the treatment of topic infections caused by antibiotic-resistant bacteria such as *P. aeruginosa*.

## Introduction

Metallic nanoparticle possess remarkable features, such as their manageable scale and the ability of fine tuning their composition, structure and shape (1). The desired electronic, optical and chemical properties of nanoparticles can be obtained through modern synthetic techniques devised in the past decades (2). Even small variations in one of these parameters may result in new properties that are absent in the analogous bulk material. Due to its high range of controllable features, metallic nanoparticles are now used in many processes, such as catalysis (3), SERS (4), electroanalytical sensors (5), plasmonics (6), photonics (7) and others.

Silver nanoparticles (AgNPs)(2), exhibit a great potential of use in many areas, due to their relative easy and controlled synthesis (8), ranging from small and simple spheroidal AgNPs (9) to more complex geometries (10, 11) and even asymmetric shapes (12–14) with solid or hollow interiors (15–17). Historrically, silver has been extensively used as an antimicrobial agent. Many different commercial products relying on silver biocidal properties have been produced, most of them in the last 10 years (18, 19). The wide variety of AgNPs as well as the intrinsic properties of metallic silver in relation to biological system and biomolecules (20) fostered the extensive use of AgNPs in medical applications, such as cancer treatment (21–23), drug delivery (24, 25), imaging (26, 27) and antimicrobial treatment (28, 29).

*P. aeruginosa* is a ubiquitous Gram-negative γ-proteobacterium and an opportunistic pathogen associated with nosocomial infections. It also affects patients afflicted by chronic respiratory diseases such as cystic fibrosis (30, 31). The relatively large genome and genetic complexity of *P. aeruginosa* (32, 33) supports its versatility and ability to colonize the soil, water bodies, as well as plants and animal tissues (33). *P. aeruginosa* infections are frequently refractoty to quinolone, aminoglycosides and β-lactam antibiotics (34). In fact, owe to its outer membrane low permeability, expression of efflux pumps, and biofilm formation, *P. aeruginosa* is one of the most difficult organisms to treat with antibiotics. Skin infections by *P. aeruginosa* on burn victims are particularly problematic, in part due to non-efficient treatments (35–37). The World Health Organization (WHO) included carbapenem resistant *P. aeruginosa* in a list of critical priority bacteria to which the research and development of new antibiotics and alternative treatments is of paramount importance (38). Although anti-bacterial treatments with visible light, such as blue light therapy (BLT) have been emerging as a non-invasive treatment for bacterial infections, the interaction of visible light with AgNPs in the range of LSPR and its possible effect on microbes has not yet been studied (39). Inspired by these findings, in the present work we use the concept of LSPR in order to enhance the antimicrobial properties of AgNPs against *P. aeruginosa* by illuminating the system with a white LED lamp. This strategy revealed a quicker and more effective treatment against *P. aeruginosa*, being a promising improvement in the fight against this pathogen.

## Results and Discussion

### AgNPs synthesis

Citrate covered AgNPs have been widely used in biological applications. Citrate, a natural anti-oxidant is not harmful to cells and is used as a carbon source by some bacteria. We synthesized citrate-covered AgNPs with a diameter of 20 ± 3 nm (Figure 1a). The extinction spectrum displayed a full width at half maximum (FWHM) of 131 nm and a maximum at 436 nm, a wavelength that matches the maximum emission from the LED floodlight used here (Figure 1b). The extinction band is broadened due to the standard deviation in particle size distribution, which represents ca. 15% of the average diameter. This is a typical spectrum for AgNPs synthesized by the citrate method (40, 41). In order to optimally observe the effects of the LSPR phenomenon, the excitation of AgNPs with an external radiation source should be performed in the region of maximum absorption. Therefore, a floodlight with an emission spectrum that superposes the AgNPs extinction spectrum was utilized (Figure 1b). To avoid the mutagenic and carcinogenic effects of UV light, a light source that did not emit in the UV region was chosen.

**Figure 1.**
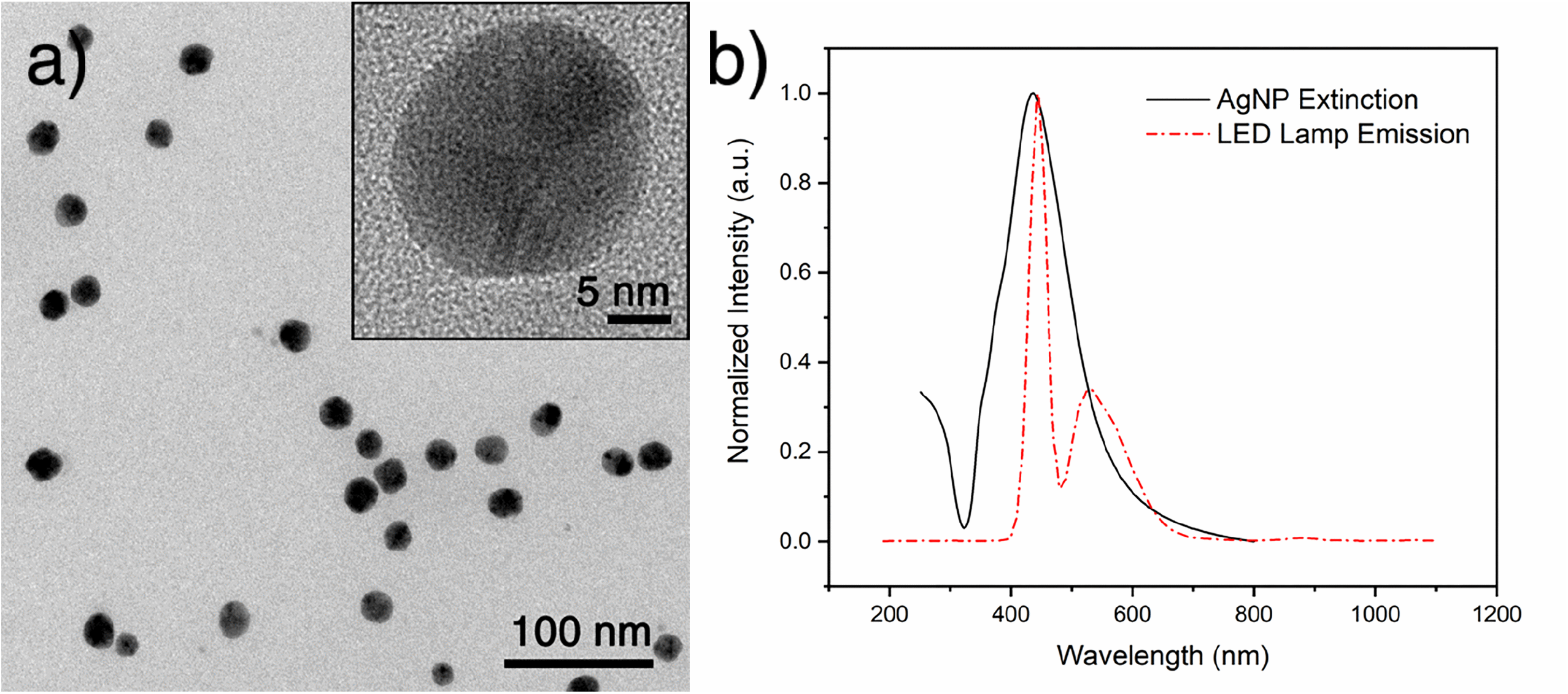
Characterization of AgNPs. (a) Representative TEM and HRTEM (inset) micrographs of citrate-coated AgNPs with an average diameter of 20 ± 3 nm. (b) Extinction spectrum of AgNPs showing the SPR peak at 436 nm superposed with the LED floodlight emission spectrum. The floodlight emission has a maximum intensity at 448 nm, with no measurable emission from 190 to 406 nm.

### Bactericidal effect of AgNPs on *P. aeruginosa*

Preliminary assays showed that the synthesized AgNPs displayed antimicrobial activity against *P. aeruginosa.* The minimal inhibitory concentration (MIC) of AgNPs was 10 µg/mL, which is comparable to values reported by similar studies for the same bacterium(42–44). However, when the AgNPs were exposed to light the MIC dropped to 5 µg/mL. The minimal bactericidal concentrations (MBC) were 20 µg/mL and 10 µg/mL under dark and light conditions, respectively. In addition, a Kirby-Bauer assay using filters containing different concentrations of AgNPs was performed in the presence and absence of light (Fig. 2). In the absence of light (Fig. 2A), an inhibition halo became apparent with AgNPs concentrations higher than 15 µg/mL and larger halos were observed with AgNPs concentrations up to 30 µg/mL. In contrast, when the AgNPs were exposed to light, 10 µg/mL sufficed to produce a halo similar to that observed when the bacteria were treated with 30 µg/mL AgNPs in the dark, suggesting that light enhances the bactericidal effect of silver nanoparticles (Figure 2B).

**Figure 2.**
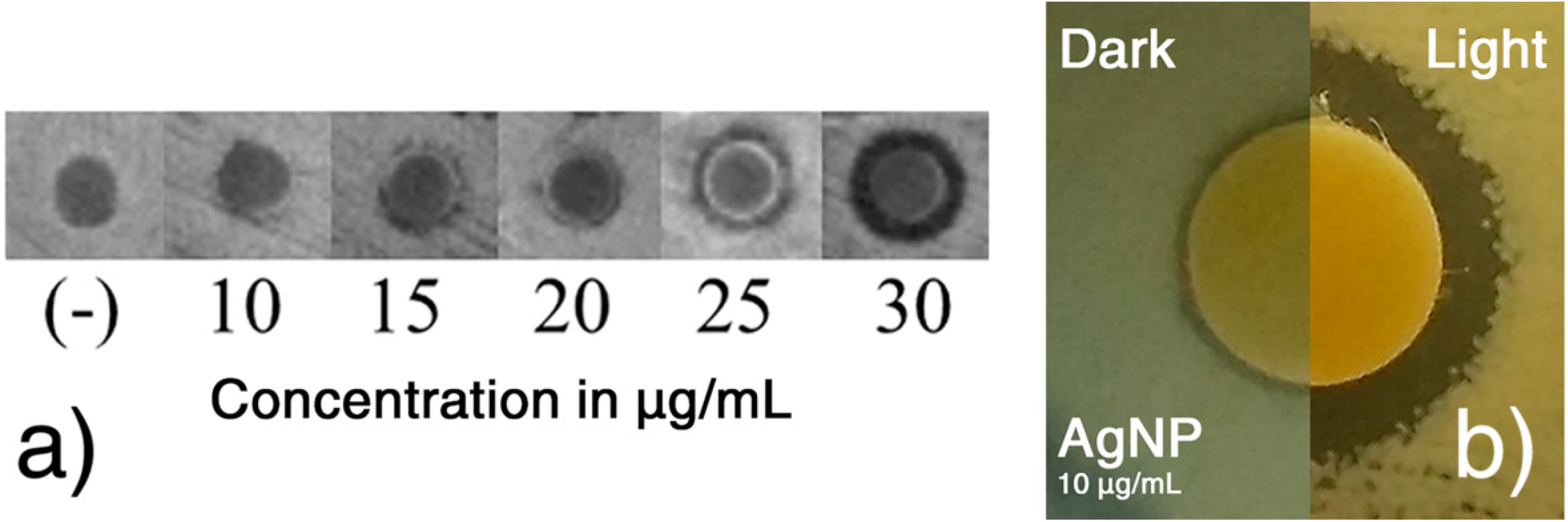
Effect of AgNPs on *P. aeruginosa* survival. (a) Disc diffusion assay: Paper discs containing each 10 µL of 0, 10, 15, 20, 25 or 30 µg/mL AgNPs were placed on Mueller-Hinton plates spread with bacteria and incubated overnight under dark conditions. (b) Effect of light on the bactericidal effect of AgNPs. Discs containing 10 µg/mL AgNPs were placed over *P. aeruginosa* lawns in the absence (left side) or presence (right side) of light.

It is worth noticing that upon light exposition, the greenish color of *P. aeruginosa* colonies, caused by the synthesis of the pigments pyoverdine and pyocyanin visually decrease. These molecules are well-known virulence factors of *P. aeruginosa* are targets for the screening of antimicrobial compounds (45–47). Accordingly, some studies have shown that light reduces the concentration of pyocyanin in *P. aeruginosa* and that this effect was dependent on light intensity and wavelength (48). It has also been shown that high doses of blue light treatment inhibited the activity of pyocyanin, staphylolysin, pseudolysin and other proteases (37, 39). Furthermore, it has been demonstrated that pyocyanin confers on *P. aeruginosa* resistance to ionic silver (49). It is thus possible that pyocyanin inhibition by LED floodlight contributes to the bactericidal effect of AgNPs on *P. aeruginosa* and diminishes the virulence of the surviving bacteria.

To further explore the effect of light exposition on *P. aeruginosa*, bacteria were grown for 60 minutes in MH medium without AgNPs in the presence or absence of light (Fig. 6). The calculated growth rates were 1.26 h^−1^ and 1.04 h^−1^ under dark and light conditions, respectively, a 17.5% reduction, suggesting that light mildly affects bacterial growth.

Next, we measured the effect of AgNPs on cell viability at different time intervals. Bacteria treated for 15, 30 and 60 minutes with 1X MIC AgNPs showed a survival rate of 12.8%, 4.4% and 1.6% in the dark and 4.5%, 0.7% and 0.2% under light conditions, respectively (Fig. 3a), while bacteria treated with 2X MIC showed, respectively, a survival rate of 20.6%, 6.4% and 2.2% (dark conditons) and 3.6%, 0.4% and 0.0% (light conditions) at 15, 30 and 60 minutes (Fig. 3b). An 1 h treatment with 10 or 20 µg/mL AgNPs under light exposition resulted in 99.8% and 100% cell death, respectively. These results corroborate the bactericidal effect of AgNPs on *P. aeruginosa* and the enhancing effect of light exposure upon AgNPs treatment.

**Figure 3.**
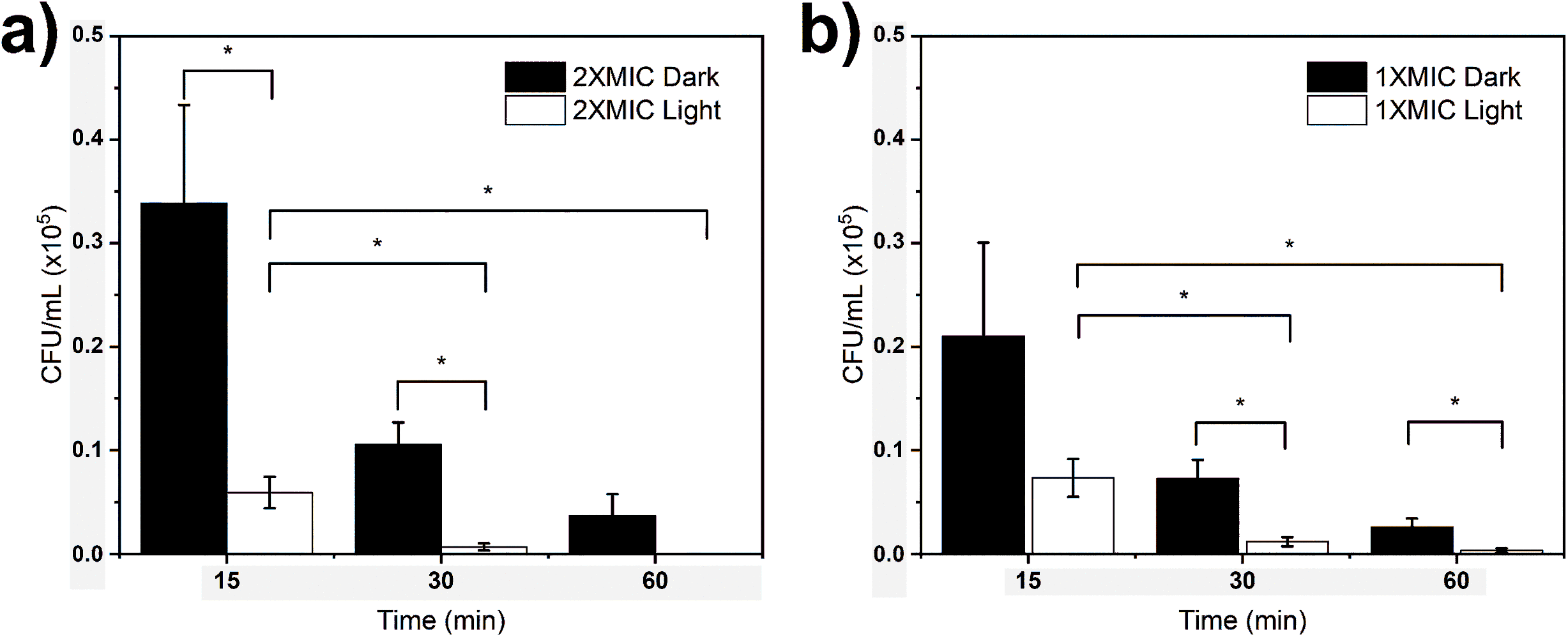
Combined effect of light and AgNPs on bacterial viability. Bacteria were grown in MH medium containing 1X MIC AgNPs (a), or 2X MIC AgNPs (b), in the absence (black) or presence (white) of light. Aliquots of 100 µL were withdrawn at 0, 15, 30 and 60 minutes to determine bacterial survival. The initial number of cells in all experiments was 1.64×10^5^ CFU/mL. Each bar represents the mean ± SD of 12 independent cultures. *p< 0.05

### Effect of light and AgNPs on ROS generation

The antimicrobial mechanism of action of AgNPs is still not fully understood mainly due to the difficulty in evaluating the contributions of nanosized metallic silver and silver ions that are released by the former (50, 51). While some studies support the idea that silver ions play the main role in cellular damage (52), others claim that the strong antibacterial activity of AgNPs is associated with the introduction of nicks in the cytoplasmic membrane which ultimately lead to cell death (53). In addition, AgNPs (and silver ions) induce the formation of reactive oxygen species (ROS) (54), which damage the cell cytoskeleton, and oxidize proteins and nucleic acids, potentially leading to chromosomal aberrations and cell demise. Also, both ionic and nanosized silver can interact with sulphur-containing macromolecules, such as proteins, resulting in their inactivation (55). Lately, a Trojan-horse like mechanism has been proposed, in which AgNPs first enter the cell and only then release silver ions (56). Previous studies showed that AgNPs promote the induction of reactive oxygen species (ROS) (57, 58) and enhance the expression of superoxide dismutase, catalase and peroxidase (58). These authors concluded that the main mechanism of antimicrobial action of AgNPs involves the disequilibrium of oxidation and antioxidation processes and the inability to eliminate ROS (58). High concentration of ROS in the bacterium lead to oxidative stress (59, 60). To test whether AgNPs causes oxidative stress, the intracellular level of ROS in *P. aeruginosa* was assessed. Bacteria were treated with 1X MIC AgNPs under light or dark conditions followed by exposure to 2′,7′-Dichlorofluorescin diacetate (DCFH-DA). This compound diffuses into the cell where it is hydrolyzed by intracellular esterases (DCFH), resulting in the production of 2’,7’-dichlorofluorescein (DCF), which fluoresces upon oxidation. The intracellular ROS level thus correlates with the level of fluorescence emitted by DCF in the presence of ROS such as H_2_O_2_, ROO^−^ and ONOO^−^ (61). Fig. 4 shows that after 1 hour in the presence of AgNPs, the level of ROS increased by 2.5-fold (dark) and 4.8-fold (light) when compared to the respective controls.

**Figure 4.**
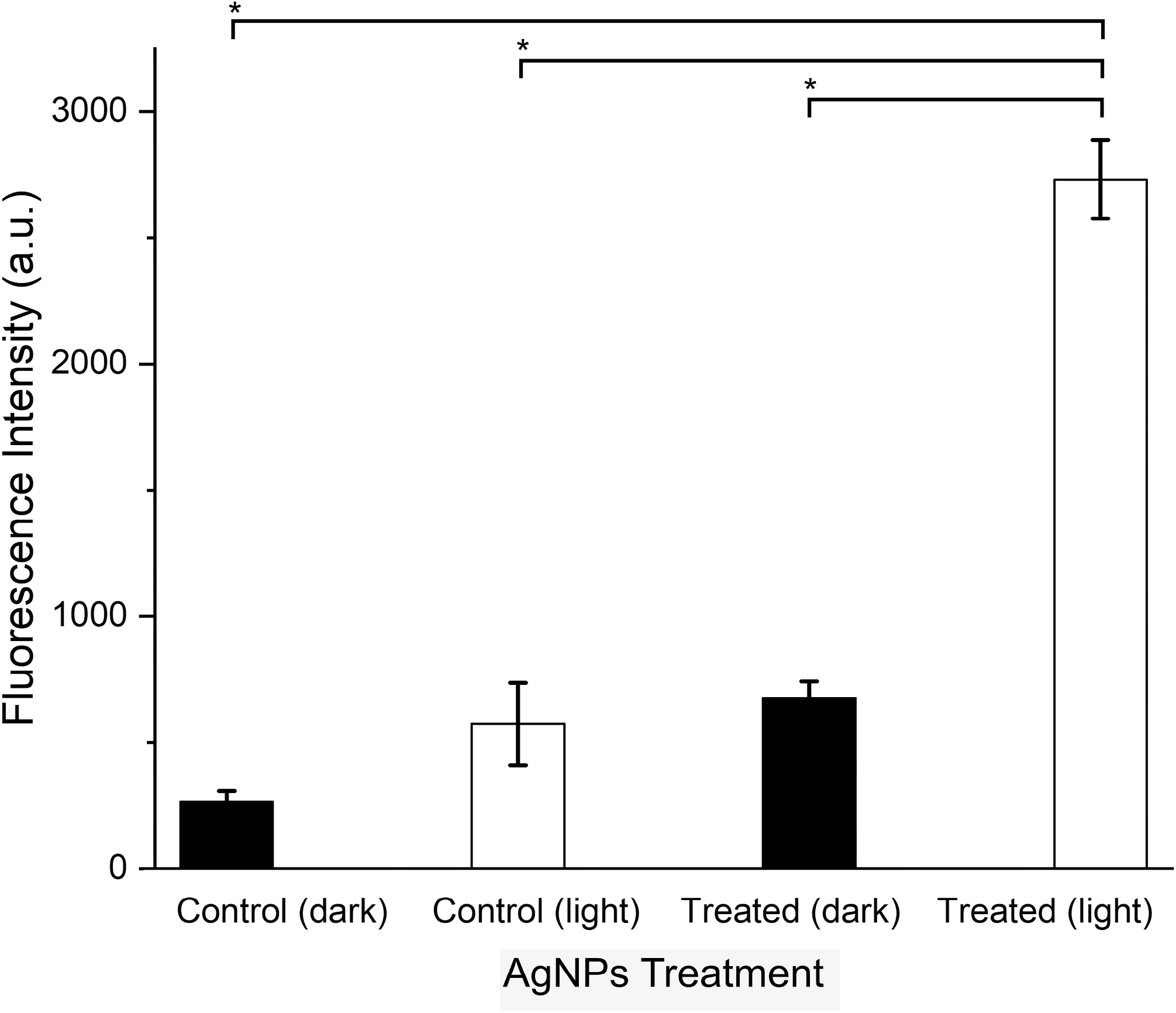
Light increases intracellular ROS generation in *P. aeruginosa* treated with AgNPs. Bacteria were exposed for 1 hour to 10 µg/mL AgNPs in the absence or presence of light. Fluorescence intensities of DCFH-DA were measured by flow cytometry. Each point represents the mean ± SD of 3 independent cultures. * denotes statistical significance with p< 0.05.

In a conducting medium, surface electrons of nanoparticles may oscillate in a resonant condition with a coupled electromagnetic field, resulting in a phenomenon called Local Surface Plasmon Resonance (LSPR), which confer on nanoparticles physical unique properties. Among them, a local temperature increase, enhancement of optical near fields surrounding the particle and the generation of hot carriers can be highlighted (62). The last one received a special attention in the field of catalysis due to the possibility of electronic or vibrational activation of substrates or reaction intermediates. Interestingly, recent studies have demonstrated the formation of ROS at the surface of metallic nanoparticles (63–65). For example, LSPR-based antibacterial treatment with gold nanoparticles has been reported (66–68).

It has been suggested that the formation of ROS by AgNPs occurs either at the particle surface or it is indirectly caused by Ag^+^ ions released from aging nanoparticles (69, 70). In addition, the light stimulus may contribute to the amount of ROS generated through a mechanism of plasmon-produced hot electrons and their interaction with water and oxygen dissolved in the medium (63–65). To assess whether exposure to light affects the production of Ag^+^ from AgNPs by an aging process, a closed dialysis membrane filled with an AgNPs suspension was submerged in distilled water and the concentration of released Ag^+^ to the solution was followed during 24 h (Fig. 5). At 0 h the concentration of silver ions was below the limit of detection of ICP-OES. The highest concentration of Ag^+^ under dark conditions was four times lower than that observed under light exposition. Although the concentration of Ag^+^ increased with light treatment, the highest observed concentration (at 6 h, under light exposition) represented only 0.4% (w/w) of the total amount of silver in the sample. Moreover, the release of Ag^+^ occurred several hours after AgNPs have already killed the bacteria (see Fig. 3). At the time in which the bactericidal effect and ROS were evaluated (1 h), it is noticeable that the measured concentrations of Ag^+^ under light and dark conditions are statistically indistinguishable. Therefore, the release of Ag^+^ is unlikely to be involved in the mechanism through which AgNPs cause bacterial death.

**Figure 5.**
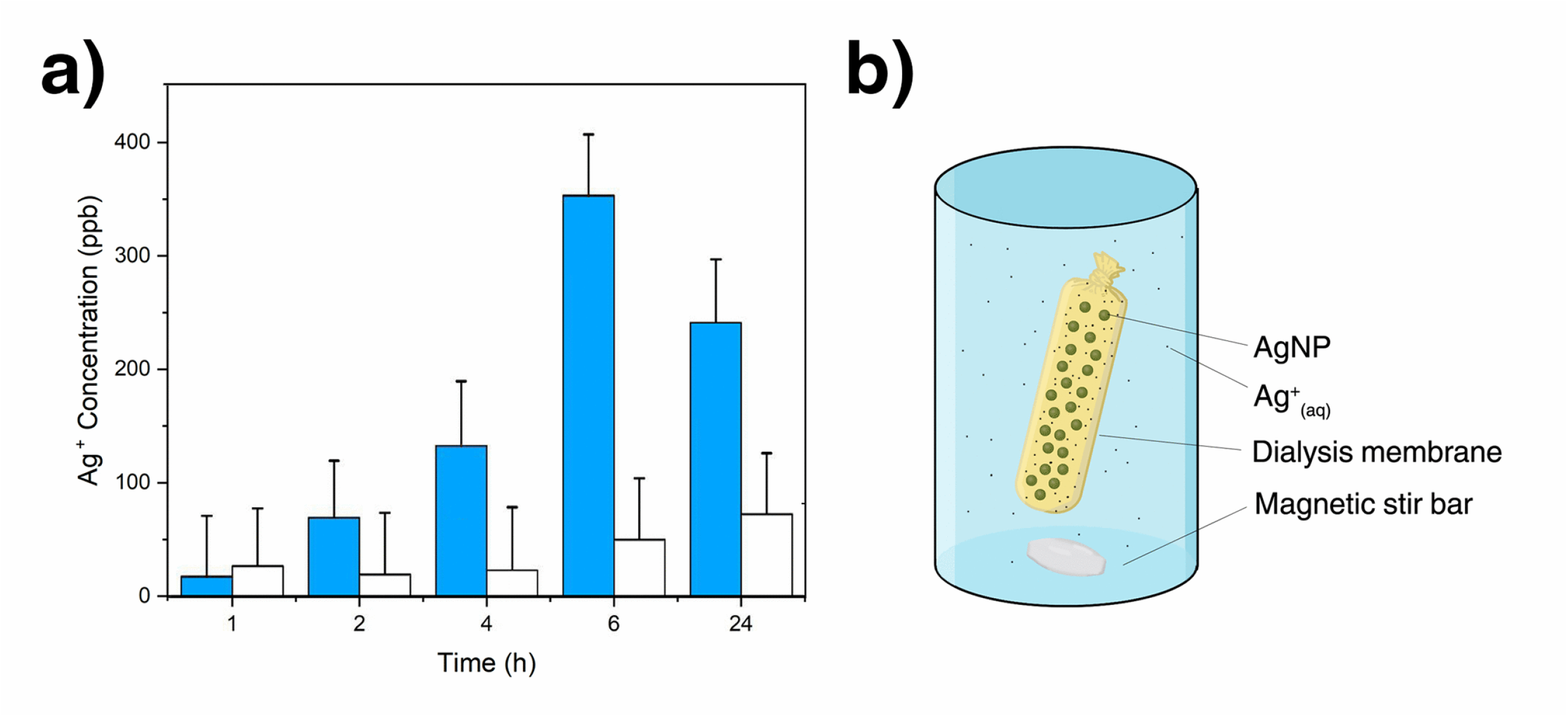
Concentration of silver ions released by AgNPs through a dialysis membrane over time in the presence (blue color) or absence of light (white color), respectively. A known amount of AgNPs was added to a tightly closed dialysis membrane and the system was submerged in distilled water under magnetic stirring. (a) Analysis of Ag^+^ content by ICP-OES. (b) Schematic representation of the dialysis method used to quantify silver ions.

**Figure 6.**
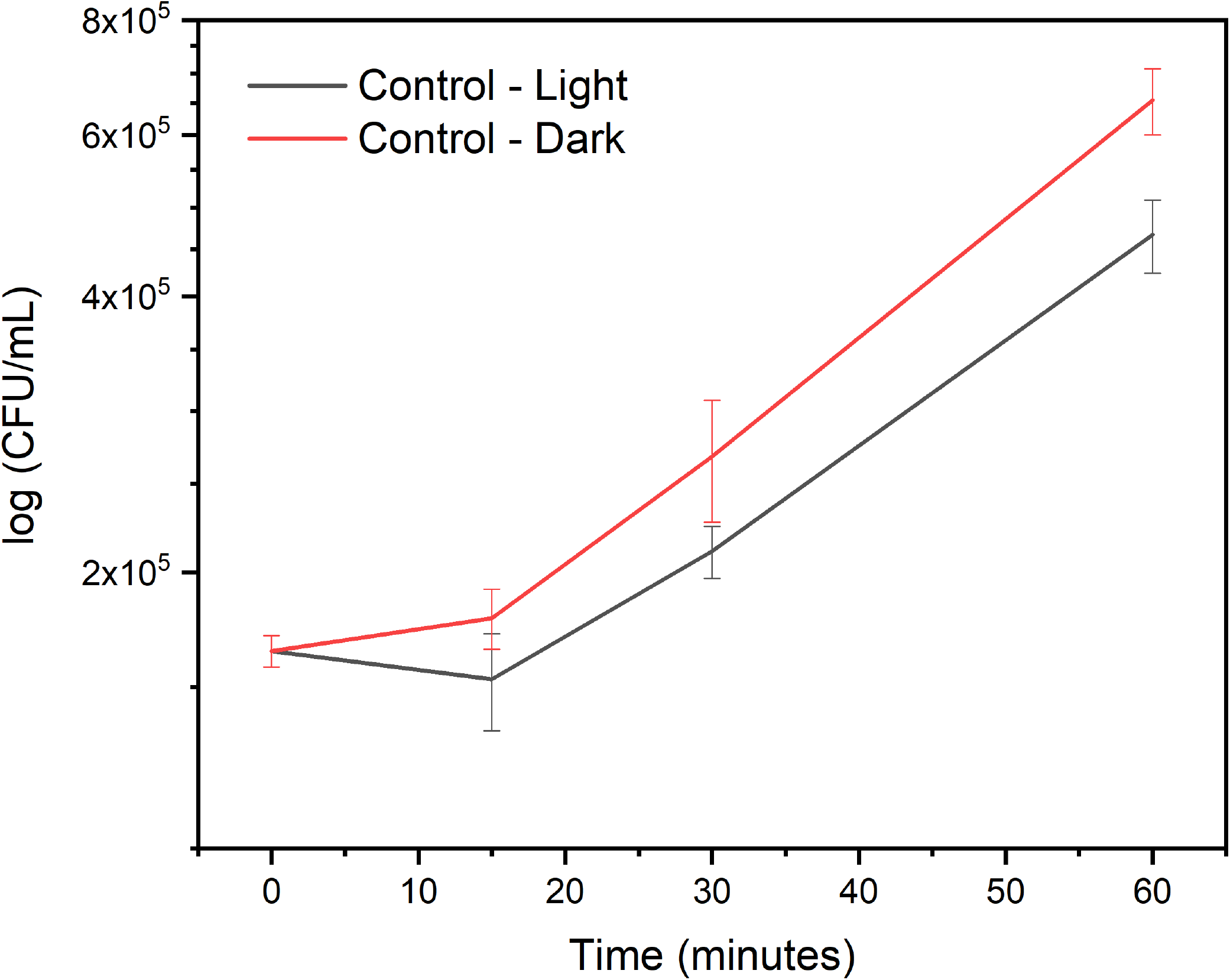
Effect of light on bacteria growing in Mueller-Hinton medium. Bacterial growth rate was determined by growing cells under dark or light conditions in MH medium and measuring the number of colony forming units (CFU/mL) at different time points.

## Conclusion

AgNPs were shown to be effective against *P. aeruginosa*. The bactericidal effect of AgNPs were intensified when the system was irradiated by light, indicating that the combination of light and AgNPs ensures a quick and more effective treatment against this pathogen. A treatment period of one hour was sufficient to cause complete cell death. These effects were associated with an increase in intracellular ROS possibly generated by the hot carrier transfer of LSPR on the AgNPs surface. AgNPs aging and the consequent release of silver ions occurred only after long periods of exposition to light. Our data illustrate a viable treatment that combines two pre-existing strategies: the use of both AgNPs and light treatment, that synergistically enhance the treatment of *P. aeruginosa* topical infections.

## Materials and Methods

### Materials

Silver nitrate (AgNO_3_, Sigma-Aldrich, ≥99%), sodium citrate (Sigma-Aldrich, ≥99%), lysogeny broth medium (KASVI, Curitiba, Brazil), phosphate buffer saline (PBS, pH 7.4), bacteriological agar (KASVI, Curitiba, Brazil), Mueller-Hinton broth (BD Difco) and 2′,7′-dichlorofluorescin diacetate (DCFH-DA, ≥97%, Sigma-Aldrich) were of analytical grade and used as received. All solutions were prepared using deionized water (18.2 MΩ) unless stated otherwise. Sterile filter paper discs of 6 mm, 96-well flat bottom plates (Corning) and 24-well flat bottom plates (Corning) were used as received.

### Synthesis of Silver Nanoparticles (AgNPs)

Preparation of AgNPs followed a slightly modified method of Lee and Meisel (71). AgNO_3_ (90 mg) was dissolved in 500 mL of distilled water. The solution was heated until boiling under magnetic stirring. Ten mL of a 34 mM sodium citrate solution were then added and the mixing which proceeded for another 1 hour. The suspension was transferred to an amber flask and stored at room temperature until further use.

### Sample preparation for electron microscopy (SEM and TEM)

One mL of the as synthesized AgNPs suspension was washed three times by successive rounds of centrifugation at 17000 rpm. The pellet containing the AgNPs was resuspended in 20 μL or 300 μL of water. For the SEM analysis, 1 μL of the concentrated suspension was deposited onto a 1 cm^2^ silicon wafer and let dry at room temperature. For the TEM analysis, 10 μL of the suspension was transferred to a standard formvar coated copper TEM grid and dried at room temperature. Particles size were analysed with the help of the ImageJ 2.0 software.

### Media and bacterial growth conditions

*Pseudomonas aeruginosa* (strain PA14) was typically cultivated in LB medium at 37°C for 16 h. The culture was then diluted 1/100 in 5 mL of MH, and grown until the exponential growth phase (OD_600_ = 0.1). Mueller-Hinton agar plates (MHA-DIFCO) were used in the bacterial viability assays in solid medium. Bacterial growth rate was determined by growing cells under dark or light conditions in MH medium and measuring the number of colony forming units (CFU/mL) at different time points.

### Determination of the minimum inhibitory concentration (MIC)

MIC determination was performed by the broth microdilution technique (Clinical and Laboratory Standards Institute – CLSI, 2015). Briefly, 10^5^ bacteria/ml were inoculated in 96-well plates containing Mueller-Hinton medium and incubated in the presence of increasing concentrations of AgNPs (from 0 to 80 μg/mL). One plate was covered with aluminium foil (dark conditions) and the other was placed on the top of it (light conditions). The plate on the top was positioned 10 cm away from a LED floodlight (50 W). Both plates were incubated overnight at 30°C. On the next day the turbidity of the cultures was measured at 600 nm in an Epoch^TM^ Microplate Spectrophotometer (BioTek).

### Antimicrobial activity of AgNPs (disc diffusion method)

The antimicrobial activity of AgNPs in solid medium was evaluated through the Kirby-Bauer disc diffusion method according to the guidelines provided (Clinical and Laboratory Standards Institute – CLSI, 2015). Mueller-Hinton agar plates were evenly swabbed with 100 µL of a PA14 culture (OD_600_= 0.1). Sterile filter paper discs (6 mm diameter, Oxoid) were uniformly placed on the plate surface and 10 µL of different concentrations of AgNPs were dropped onto each disc. The negative control was sodium citrate at the same concentration of the AgNPs suspension. Plates were incubated overnight at 30 ºC and the halos formed by antibiotic activity against the bacteria were measured with the help of the ImageJ 2.0 software.

### Kinetics of AgNPs antibacterial effect

Two 24-well plates were filled with 1.5 mL of a mixture of MH medium and AgNPs (0, 1X and 2X MIC). A final concentration of 10^5^ cells/mL at the exponential growth phase was added to each well. One of the plates was covered with aluminium foil (dark conditions) and the other one was placed on its top (light conditions). Both plates were statically incubated at 30ºC, while the plate on the top was 10 cm away from a LED floodlight (50 W). Aliquots of 100 µL were withdrawn at 0, 15, 30 and 60 minutes from both plates and subsequently diluted in 0.9% NaCl, plated on LB-Agar and incubated for 24 h at 37 ºC. The number of colony forming units (CFU/mL) was determined. This experiment was performed with twelve biological replicates.

### Intracellular ROS assay

Three independent overnight cultures of *P. aeruginosa* were diluted 1:100 in fresh MH medium, incubated under agitation until reaching the concentration of 10^8^ cells/mL. Different concentrations of AgNPs and 20 µM DCFH-DA were then added. The cultures were incubated at 30 ºC for 1h in the presence and absence of light exposition. Then, 1 mL of each culture was withdrawn and centrifuged at 8000 rpm for 5 minutes. The pellets were washed with 1 mL of PBS and suspended again in 100 μL of PBS. The fluorescence emitted by the intracellular oxidation of the dye was determined using the BD Accuri^TM^ C6 flow cytometer at the excitation wavelength of 488 nm and at an emission wavelength of 535 nm.

### Quantification of released Ag^+^

Six dialysis membranes each filled with 20 mL AgNPs were placed in 800 mL beakers filled with 500 mL distilled water under magnetic stirring. Three of them were placed 10 cm away from the floodlight, while the other three were covered in aluminium foil. After 1, 2, 4, 6 and 24 h, 5 mL aliquots of water from each beaker were withdrawn and stored for further analysis. Silver ions were detected by iCAP^TM^ 7000 Plus Series ICP-OES (Thermo Scientific^TM^).

### Statistical

Statistical significance was calculated using the analysis of variance (ANOVA) followed by Tukey pairwise comparison. Values of p ≤ 0.05 were considered statistically significant.

## Acknowledgments

This work was supported by FAPESP (grant number 2018/13492-2; 2015/26308-7). E.Y.V was funded by a postdoctoral fellowship from PNPD/CAPES (grant 1680341). R.T.P.S was funded by a PhD fellowship from CAPES (grant 88882.328241/2019-01).

